# Sleep stages antagonistically modulate reactivation drift

**DOI:** 10.1101/2023.10.13.562165

**Authors:** Lars Bollmann, Peter Baracskay, Federico Stella, Jozsef Csicsvari

**Affiliations:** IST Austria, Klosterneuburg, Austria; Donders Institute, Nijmegen, The Netherlands

**Keywords:** Hippocampus, Sleep, Sharp-Wave Ripples, REM, Place Cells, Reactivations, Replay, Learning, Hidden Markov Model, Bayesian Decoding

## Abstract

Hippocampal reactivation of waking neuronal assemblies in sleep is a key initial step of systems consolidation. Nevertheless, it is unclear whether reactivated assemblies are static or whether they reorganize gradually over prolonged sleep. Here, we tracked reactivated CA1 assembly patterns over ∼20 hours of sleep/rest periods and related them to assemblies seen before or after in a spatial learning paradigm. We found that reactivated assembly patterns were gradually transformed and started to resemble those seen in the subsequent recall session. Periods of rapid eye movement (REM) sleep and non-REM (NREM) had antagonistic roles: while NREM accelerated the assembly drift, REM countered it. Moreover, only a subset of rate-changing pyramidal cells contributed to the drift, while stable firing rate cells maintained unaltered reactivation patterns. Our data suggest that prolonged sleep promotes the spontaneous reorganization of spatial assemblies, which can contribute to daily cognitive map changes or encoding new learning situations.

## Introduction

It has been long established that sleep promotes the recall of previously acquired memories, and there is ample evidence linking sleep to systems consolidation ^1^. Furthermore, in many instances, the hippocampus plays a critical role in sleep-associated memory stabilization ^2^. A long-standing idea suggests that the hippocampus acts as an intermediate storage area for recently acquired memories, and such memories are reactivated in sleep ^3,4^. Subsequently, the reactivation of these memory traces would promote their transfer to other cortical areas for consolidation and long-term storage ^5,6^. This hypothesis has received consistent experimental support from a multitude of human and animal studies ^7^. Amongst these, rodent electrophysiological experiments could directly examine the content of neuronal activity during reactivation and test their relationship to memory recall. These showed that the rate of reactivation is increased after the experience of novel learning situations, and the content of reactivation predicts subsequent memory performance ^8–11^. Furthermore, suppressing or prolonging sharp-wave ripples (SWR), a local field potential event that usually accompanies hippocampal reactivation, disrupts or facilitates, respectively, subsequent memory recall ^9,12^. Finally, content-specific disruption of reactivation events leads to selective impairments in the recall of the disrupted memory ^13^.

So far, hippocampal reactivation of neuronal activity has been primarily studied during brief (<1 h) periods of sleep or during waking behavior, with the latter linked to different functions such as decision making or short-term memory. These works may have only examined the initial stages of a more complex sleep reactivation process, which may continue for prolonged periods ^14^. Reactivation events can be detected over longer (4-6h) periods of time ^15,16^ and in these studies, neuronal patterns of recent experiences were preferentially reactivated during the entire sleep duration. However, it is not clear whether the neural signature of specific experiences remains unchanged over extended periods or, eventually, it undergoes some form of modification and rearrangement due to early consolidation or other network processes.

Here, we quantified reactivation dynamics during up to 20 hours of quiet rest periods and sleep following spatial learning and tested to what degree reactivated patterns represented previous spatial learning patterns or resembled those seen at subsequent memory retrieval. We provide evidence for the transformation of reactivated memory traces and show that many of the transformed reactivated neuronal patterns will recur in the subsequent spatial memory recall trials. We also show that REM and non-REM periods exert opposite actions on this transformation. Finally, we show that this transformation is driven by a subgroup of pyramidal cells only and that interneuron activity also reflects the speed of transformation.

## Results

We performed 128-channel wireless recordings in the dorsal CA1 region in three Long-Evans rats using bilaterally implanted 32-tetrode microdrives. Continuous recordings were performed for a period of >24 hrs, covering the learning of a novel set of goal locations on a cheeseboard maze ^8^, an interleaved, extended rest period at the home cage, and the recall of the previously learned goal locations (Figure 1a). We extracted the activity of multiple, single units of putative pyramidal cells and interneurons that exhibited stable spike features during the entire recording (Figure S1a-d). Animals learned the novel set of goal locations within the first three trials of the learning session, and they were able to directly retrieve the rewards at subsequent trials, even at the first trial of the recall session the next day (Figure 1b).

**Figure 1:**
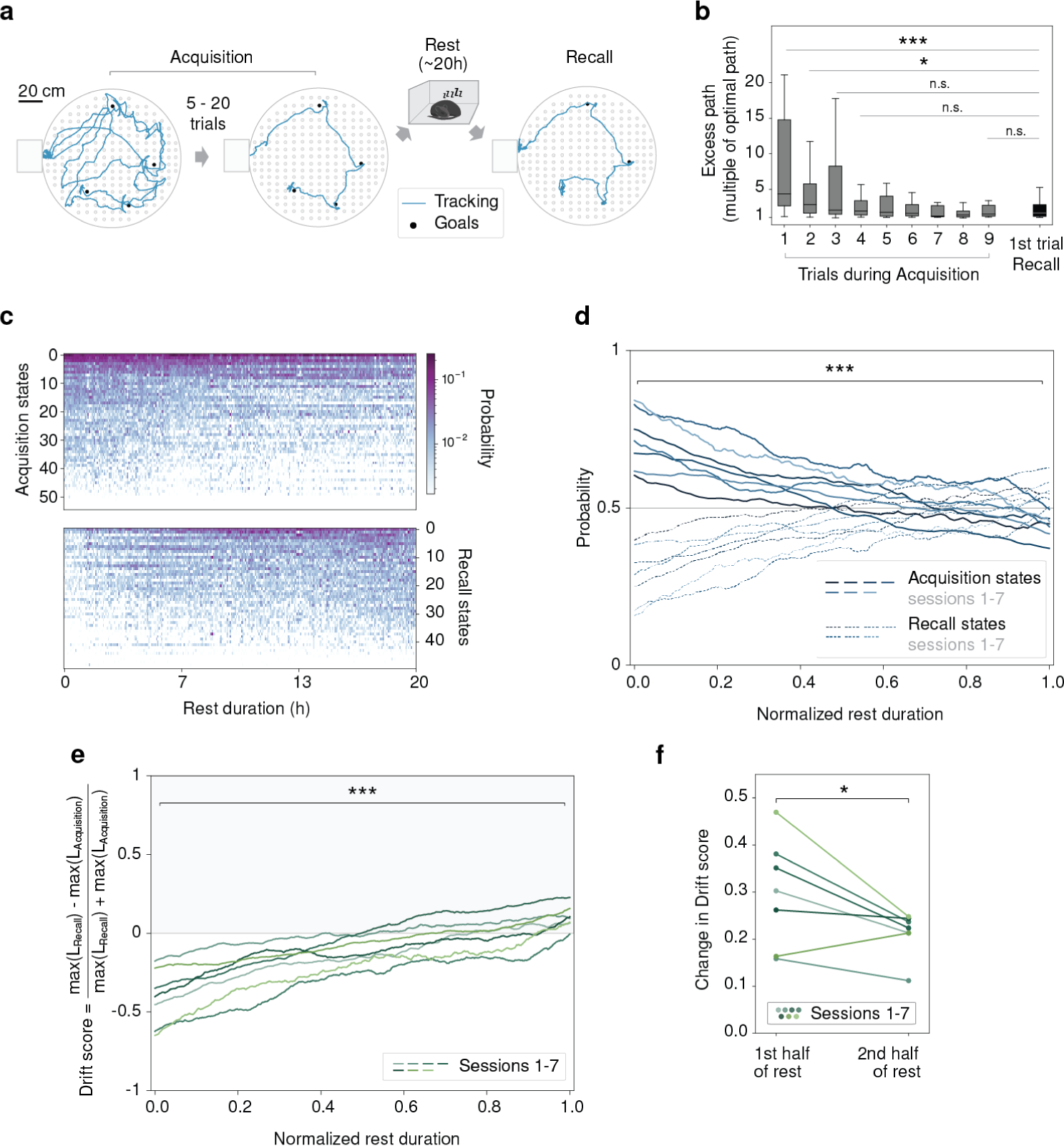
Reactivation drift during prolonged periods of sleep following spatial learning on the cheeseboard. **(a)** Behavioral sessions illustrated by animal tracking data for one example session: learning a novel set of goal locations (acquisition), long (∼20h) quiet rest/sleep period (rest) and recall of goal locations (recall). **(b)** The excess path is measured as a multiple of the optimal path to retrieve all goals during the first nine trials of memory acquisition and the first trial of recall. (* p < 0.05, *** p < 0.001, n.s. p > 0.05, one-sided Mann-Whitney U test, Bonferroni correction). **(c)** Instances when acquisition and recall pHMM states were detected with the corresponding decoding probabilities during rest for one example session. **(d)** The probability of detecting acquisition over recall pHMM states at different stages of rest for all sessions (i.e., the proportion of acquisition or recall decoding windows identified). The detection probabilities at t=0 differ significantly from the corresponding values at t=1 (p < 10e-8, one-sided t-test). **(e)** Drift score as a function of normalized sleep duration for all sessions. The drift score is below zero at t=0 (p < 0.001, one-sided t-test), above zero for t=1 (p < 0.01, one-sided t-test) and significantly different between t=0 and t=1 (p < 10e-5, one-sided t-test). **(f)** Drift score changes more strongly during the first half of sleep than during the second half of sleep (alternative hypothesis: ratio first half/second half is greater than 1, p < 0.05, one-sided t-test).

We used a Poisson Hidden Markov Model (pHMM) to identify the activity of different pyramidal cell assemblies during the learning and recall sessions (Figure S1e-i). The model did not use any prior assumption about assembly coding nor the spatial coding of place cells: it was only based on the correlated temporal firing structure of pyramidal population activity in order to identify activation states. To see whether pHMM states expressed spatial selectivity, we plotted the location of the animal at each time a specific state was active. The majority of pHMM states tended to occur only at a narrow set of locations on the cheeseboard maze, many near the goal locations, even though the states were established without prior knowledge of the location of the animal (Figure S2a-b). Consequently, the pHMM was able to decode the animal’s location with an accuracy comparable to that obtained by a standard Bayesian decoding procedure (Figure S2c-f).

Next, we used the pHMM to trace the reactivation of waking assembly patterns during the prolonged rest periods between the learning and recall sessions. For the majority of the rest period, the animal was immobile and sleeping or awake but immobile in its home cage. In examining reactivation, we only considered quiet rest periods and excluded periods when the animal moved within the home cage. In the decoding procedure, we used a temporal binning that conserved the number of spikes in any bin (12 spikes, if not stated otherwise). Within the rest period, we differentiated rapid eye movement (REM) sleep periods and the remaining quiet rest non-REM (NREM) periods and identified reactivation events during continuously detected time windows in REM sleep and in SWRs during NREM periods. Reactivations of specific patterns were identified by determining, for each fixed-spike bin, the pHMM state that matched the activity with the highest likelihood, assuming Poisson firing probabilities (Figure S1e and see Methods). To see whether reactivated neuronal patterns underwent a transformation during the prolonged rest periods, we separately compared rest patterns to the set of pHMM states of both acquisition and recall and tracked their likelihoods during rest. We found that in the first half of the rest session, acquisition pHMM states were preferentially decoded (Figure 1c-d). In contrast, later, acquisition and recall states were decoded with similar probability, or the decoding of acquisition states showed a weak preference. This indicated a drift in the reactivation patterns: activity patterns were initially similar to those expressed in the acquisition stage but progressively morphed towards those seen in the recall stage following sleep.

Next, we tracked over time how the reactivation of learning-associated neuronal patterns was gradually overtaken by recall-associated patterns. To do so, for each reactivation event, we compared the decoding log-likelihood of the learning and recall pHMMs states, and a drift score was calculated, defined by the normalized difference of the log-likelihoods (Figure S1e). A negative drift score indicates the reactivation of acquisition pHMM states, while a positive score corresponds with patterns more similar to the recall states. The drift score gradually increased over time, reaching positive values only after the second half of the rest session, after 7-8 h (Figure 1e). However, the speed of change of the drift score was faster during the first half of sleep than in the second half, suggesting that the majority of new recall states started to be active already in the first half of the rest session (Figure 1f). We obtained similar results when Bayesian position prediction probabilities were used to calculate the drift score (Figure S2g-h).

In summary, the neural activity during rest gradually changed from being very similar to the acquisition of the memory at the beginning of rest towards being more similar to the neural activity during recall at the end of rest.

We next examined whether reactivation drift exhibited differences in REM and NREM periods. When we plotted drift scores over time after performing a smoothing window averaging, we observed temporal fluctuations with a zigzagging appearance. We asked how these repeated upward and downward trends aligned with the transitions between NREM and REM phases. We found that the smoothed drift scores tended to increase during NREM while decreasing during REM periods, hence demonstrating that the fluctuations we saw were driven by the NREM-REM cycles (Figure 2a). To quantify this effect, we calculated the change in smoothed drift scores for each REM and NREM epoch. The average drift score changes were positive for NREM and negative for REM, with only about a quarter of the REM and NREM epochs showing the opposite tendencies (Figure 2b, i.e., positive REM and negative NREM). Consequently, the cumulative effect of NREM periods on the drift score was positive, whereas REM periods showed a negative cumulative effect (Figure 2c and Figure S2i). We found a difference of two orders of magnitude when we compared the cumulative change in the drift score throughout the rest with the epoch-by-epoch change in NREM and REM (Figure 2d). Short- and longer-term changes in drift score occurring during NREM and REM epochs were not correlated (Figure S2j). Hence, the drift score fluctuated strongly on shorter timescales without a clear direction, whereas the long-term change was directed and much smaller in magnitude, which implies that different processes drive the short-term fluctuation and the long-term drift.

**Figure 2:**
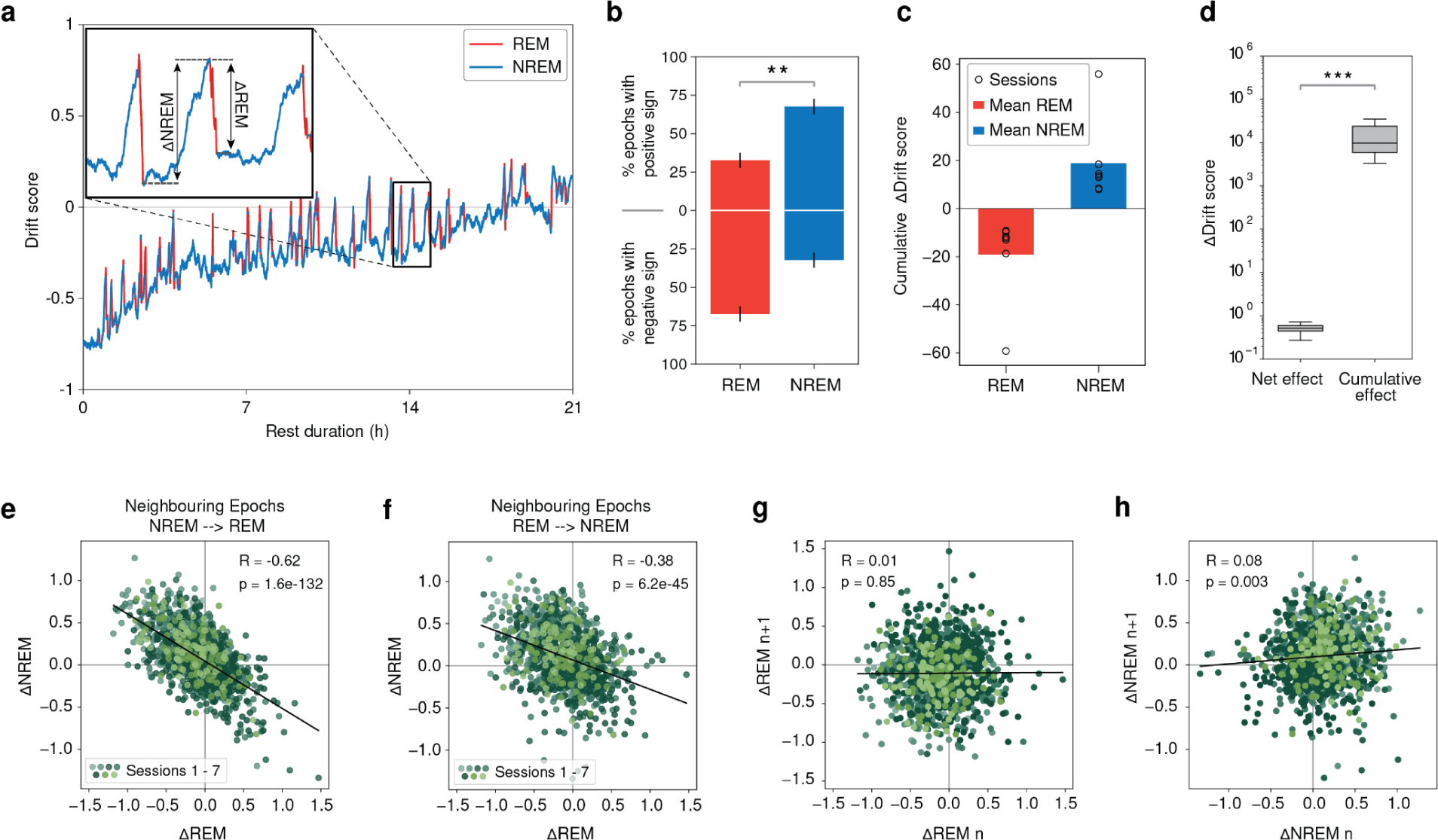
NREM periods accelerated the reactivation drift, while REM periods countered it. **(a)** Smoothed drift score as a function of sleep duration (REM periods in red, NREM periods in blue) for one example session. Inset: time expanded trace illustrating the ΔDrift score calculations **(b)** Contributions of REM and NREM epochs to drift. For each epoch, the ΔDrift score was computed as the difference in the drift score between the end and beginning. The percentage of epochs with positive and negative values are depicted for REM (red) and NREM (blue). Data from all sessions (mean±SEM, ** p < 0.001, two-sided Mann-Whitney U test). **(c)** Summed ΔDrift score for REM and NREM epochs for all sessions (p < 0.01, two-sided Mann-Whitney U test). **(d)** Net effect of change in drift score (difference in drift score between the beginning and the end of rest) and the cumulative effect of change in drift score (sum of absolute ΔDrift score for REM and NREM periods). Data from all sessions (p < 0.01, two-sided Mann-Whitney U test). **(e)** ΔDrift score values of NREM with subsequent REM epochs (R = −0.62, p = 1.6e-132). **(f)** ΔDrift score values of REM with subsequent NREM epochs (R = −0.38, p = 6.2e-45). **(g)** ΔDrift score values for subsequent REM epochs (R = 0.01, p = 0.85). **(h)** The boxplot of ΔDrift score values for subsequent NREM epochs (R = 0.08, p = 0.003).

Next, we tested whether the magnitude of up- and downward shifts in the drift scores across NREM and REM periods were independent or whether the drift in subsequent sleep epochs was interrelated. The drift score changes in neighboring NREM vs. REM periods were negatively correlated, and the NREM to REM drift score changes exhibited stronger correlations than the REM to NREM ones (p=3e-15, Fisher’s z-test, Figure 2e-f). However, the correlations of the drift score changes were weaker when subsequent REM or NREM periods were compared (Figure 2g-h). Overall, these results suggested that NREM periods drove the drift of reactivated assemblies toward the ones that will re-emerge during the recall session, while REM periods countered this effect. The correlations also suggest that the NREM drift influences the magnitude of subsequent REM drift more than REM drift influences the subsequent NREM changes. Next, we tested the contribution of individual cells to the observed reactivation drift. To do so, we characterized the firing rate changes of neurons between the acquisition and the recall period (Figure 3a and Methods). We observed that some pyramidal cells exhibited stable firing rates across the two sessions while others either increased or decreased their firing rate from one to the other (Figure 3b).

**Figure 3:**
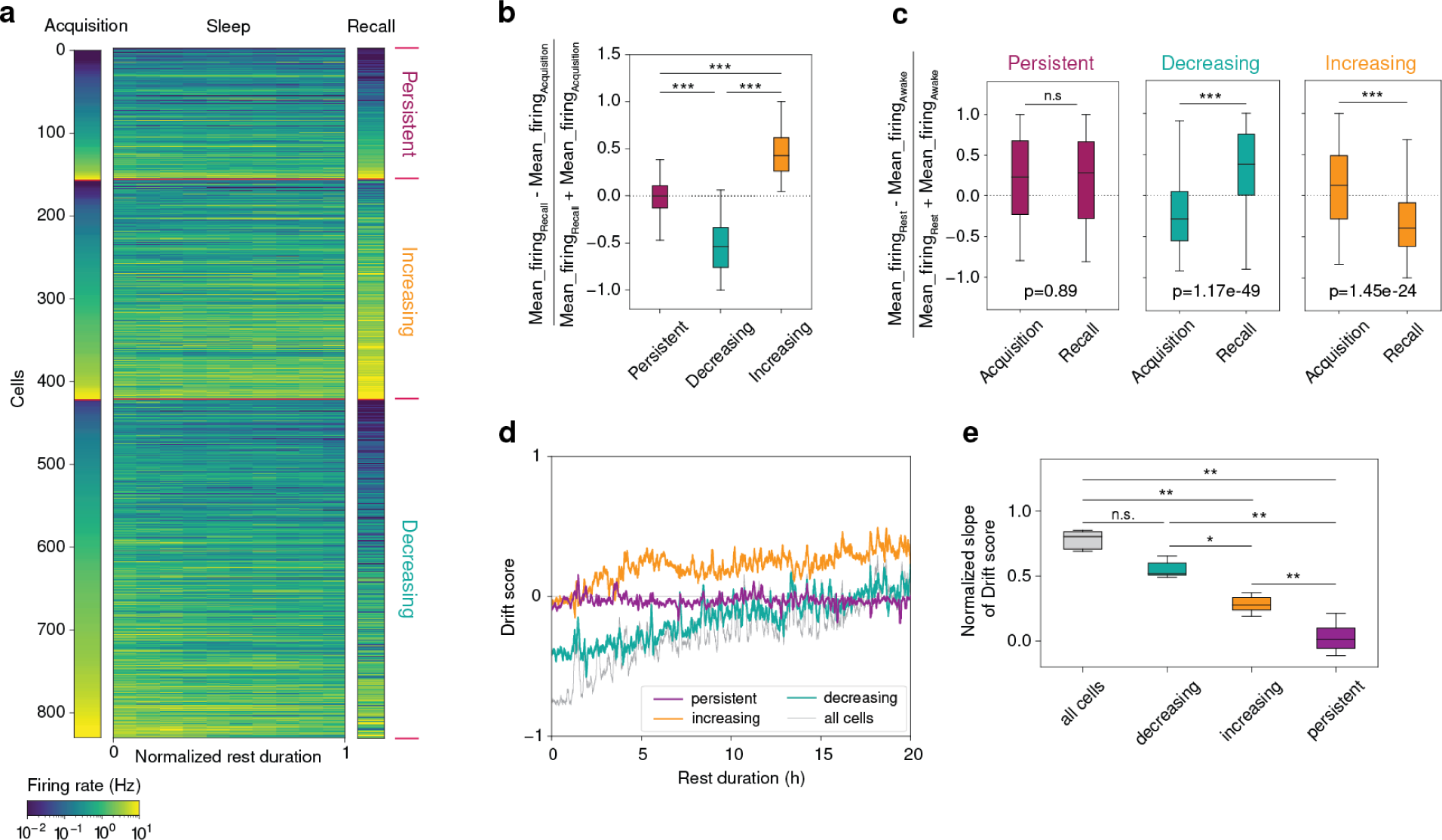
Contribution of different firing rate modulation cell groups to reactivation drift. **(a)** Average firing rates of persistent, increasing, and decreasing cells during acquisition, recall, and fixed periods of rest. Cells from all sessions are shown. Cells were classified depending on whether their firing rate distributions exhibited significant differences (p<0.05) between acquisition and recall. **(b)** Normalized firing rate change from acquisition to recall for persistent, decreasing and increasing cells (all p < 0.001, two-sided Mann-Whitney U test, Bonferroni correction). **(c)** Normalized firing rate change between rest vs. acquisition or recall (n.s. p > 0.05, *** p < 0.001, two-sided Mann-Whitney U test). **(d)** Drift score using either all cells (gray), only persistent cells (violet), only increasing cells (orange), or only decreasing cells (turquoise) for one example session. **(e)** Slope fit to drift score, calculated as the average of the slope in quadrants of rest (** p < 0.01, * p < 0.05, n.s.: p > 0.05, two-sided Mann-Whitney U test, Bonferroni correction).

When we looked at the mean rate of the cells as specified in their pHMM mean firing rate vectors (Figure S1e), the same trend appeared when acquisition and recall states were compared (Figure S3a). This showed that with the activation of recall phMM patterns later on during the rest session, cells showed more similar firing rate trends to those seen later during the recall session. More than half of the recorded cells were from the decreasing group, and only a smaller portion of cells had stable or increasing firing rates over the entire experiment (Figure S3b). Cells with persistent activity, on average, exhibited lower firing rates than the rate-changing groups but maintained a comparatively higher rate during sleep (Figure 3c and Figure S3c-e). We did not find differences in SWR firing gain or waveform stability between the different subsets (Figure S3f-g), and different cells recorded from the same electrode could exhibit different rate modulations (i.e., belong to different groups) (Figure S3h). Spatial information of persistent cells was lower compared to decreasing cells during the acquisition and lower compared to the increasing cells during recall (Figure S3i). Nevertheless, we did not observe major differences in the goal-related remapping during learning or recall between persistent and rate-changing cells (Figure S4a-d).

When we performed the reactivation drift analysis using only the persistent cells, the drift score remained stable near zero across the entire rest period, indicating a high similarity of acquisition and recall patterns and their consistent reactivation during sleep (Figure 3d-e). However, reactivations by the other two groups yielded similar drift score changes to that calculated with all cells. This result suggests that firing rate changes of the rate-changing cell groups were the primary drivers of the reactivation drift. Furthermore, we also found that the spatial coding of the environment remained more similar for the persistent cell group than for the rate-changing groups, as measured with population vector similarity across acquisition and recall (Figure S4e).

To further investigate the contribution of pyramidal rate changes to reactivation drift, we tested whether rate changes seen during individual REM and NREM epochs correlated with the magnitude of drift score changes at the same epochs: for the increasing group, we detected a positive correlation, while a negative correlation was seen for the decreasing cells (Figure 4a-g). Similarly to the drift score changes, neighboring REM and NREM periods exhibited correlated firing rate changes for both the decreasing and increasing groups, with a stronger effect for NREM periods being followed by REM periods (p=1.23e-22 for decreasing cells, p=4.91e-19 for increasing cells, Fisher’s z-test, Figure S4f-i). Firing rate changes between sleep epochs of the same type were only weakly correlated (Figure S4j-m).

**Figure 4:**
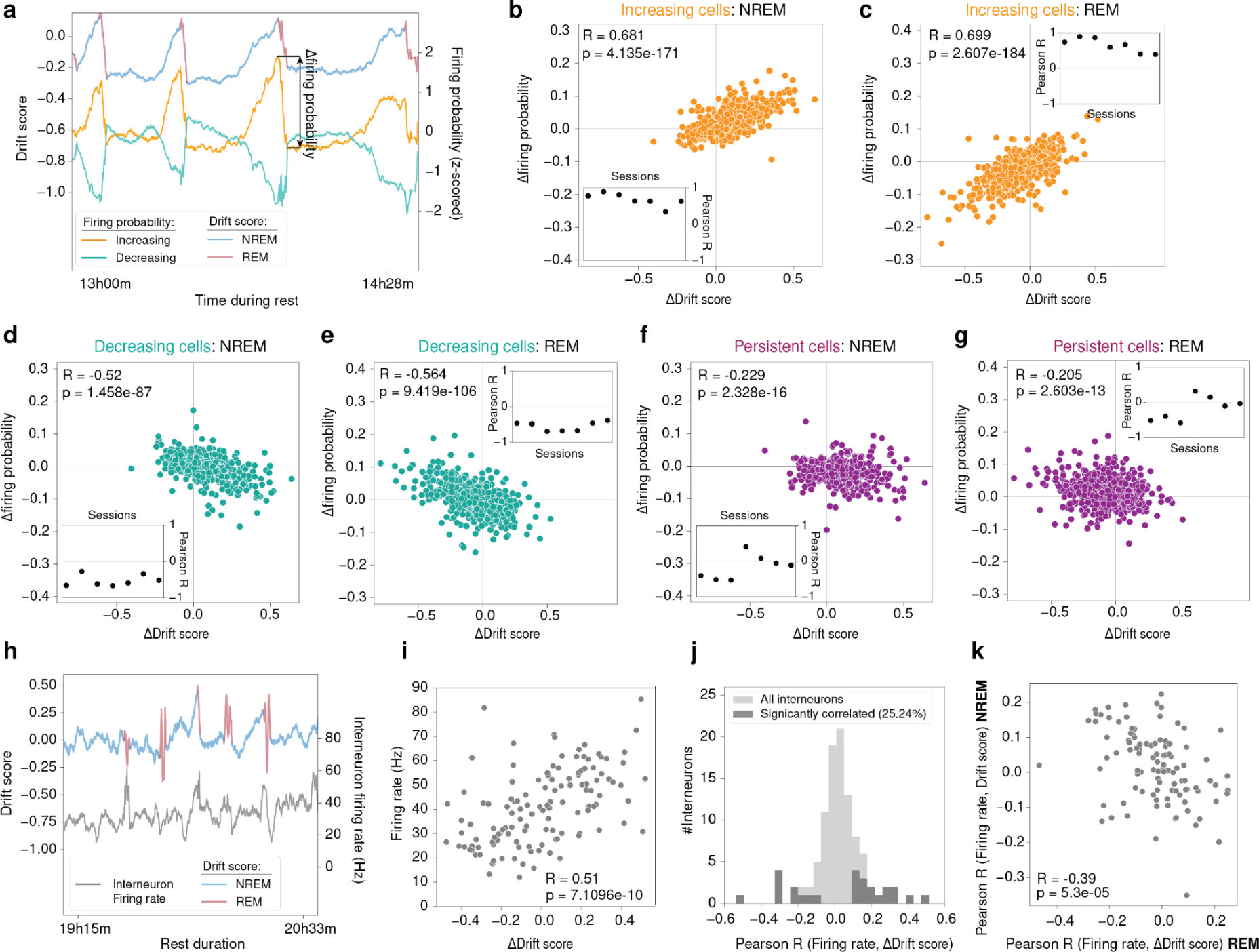
Reactivation drift correlates with the firing rate modulations of pyramidal populations and a subset of interneurons. **(a)** Illustration of drift score and firing probability fluctuations of decreasing and increasing cells for one example session. **(b-g)** Correlation between ΔDrift score and the change in firing probability for increasing cells during NREM and REM periods. Inset: correlation values per session(increasing cells: NREM **(b)** R = 0.68, p = 4.13e-171, REM **(c)** R=0.70, p = 2.61e-184; decreasing cells: NREM **(d)** R = −0.52, p = 1.46e-87, REM **(e)** R = −0.56, p = 9.42e-106); persistent cells: NREM **(f)** R = −0.23, p = 2.33e-16, REM **(g)** R = −0.21, p = 2.60e-13). **(h)** example for the correlated fluctuation of the drift score and the firing rate of an example interneuron. **(i)** Correlation of firing rate and ΔDrift score for the interneuron from (h) (R = 0.51, p = 7.11e-10). **(j)** Distribution of Pearson R values for interneuron firing rate and delta score using REM and NREM epochs. A significant proportion of interneurons exhibited a significant (p<0.05) correlation between firing rate and ΔDrift score (p<0.001, binomial test). **(k)** Pearson R values for interneuron firing rate and ΔDrift score calculated separately in NREM and REM epochs (R = −0.39, p = 5.3e-5).

Next, we tested whether the interneuron activity was related to reactivation drift. The activity of some interneurons strongly reflected the drift score changes in REM and NREM epochs (Figure 4h-i). In agreement with this, 25.2% of the interneurons exhibited a significant correlation between their firing rate and the change in drift score (Figure 4h-k). Some of these interneurons increased their firing rates with the magnitude of rate changes during both REM and NREM epochs, while others reduced it.

After separating REM and NREM epochs, a significant fraction of interneurons only maintained a relationship between mean firing and the change in drift score for REM epochs (S4n-o). However, interneurons exhibited a reversed relationship in REM and NREM sleep between mean firing rate and change in drift score: if an interneuron’s firing rate showed a positive relationship with the change in drift score in REM, the relationship was negative in NREM and vice versa. This effect modulated all recorded interneurons regardless of the strength of the firing rate to drift relationship. In summary, the firing rates of a subset of interneurons mirrored the fluctuations of the drift score. Moreover, the whole interneuron population exhibited a reversed coupling to REM and NREM periods in terms of their net firing rate modulation, with some exhibiting preferred rate increases either in REM or NREM compared to the other state.

## Discussion

Here, we demonstrated that reactivated patterns in the hippocampus reorganize during prolonged rest periods (∼20h), after spatial learning of novel goal locations in a familiar environment. In the first 8-10 hours of rest, these reactivated patterns primarily represented the previous learning session patterns, but later, they were gradually overtaken by those resembling future patterns expressed in the subsequent recall session. We also found that this reactivation drift was antagonistically modulated by the succession of NREM and REM epochs: while NREM periods favored the drift towards new patterns of the recall session, REM periods had the opposite effect. Hence, we observed the gradual reorganization of reactivated hippocampal assembly patterns, and the reorganization was regulated by NREM-REM sleep cycles.

### Reactivation drift reflects place map reorganization

Our ability to detect the progressive transformation of hippocampal reactivated patterns relied on their continuous monitoring over prolonged periods of sleep. Previous studies in which intermediate sleep durations (<6 h) were used observed a relatively stable reactivation expression ^15,16^ . In our case, although the majority of the novel recall patterns emerged in the first half of the rest periods, these replaced the prior learning patterns only gradually, and it took about 8-10h for recall assembly patterns to match or overtake the learning patterns. This can explain why previous studies of shorter sleep durations observed stable reactivation even after the prior exploration of novel environments.

However, there is experimental data, suggesting that reactivated sleep patterns do not exclusively represent recent experiences. Even early reactivation studies were able to detect a weak but significant similarity between sleep activity patterns and subsequent waking patterns of familiar environments under conditions in which the animal visited the familiar environment a day before at the earliest ^17^. This suggests that less recent experiences are also reactivated in sleep, including those the animal experienced in previous days. However, later preplay was observed in which sleep temporal patterns were similar to place cell firing patterns that emerge later in a novel environment ^18–20^. Even in familiar environments, one can observe a representational drift of place maps, where some new place cells emerge while others disappear in later exploration of the same environment ^21–23^. Similar to preplay, neuronal patterns during sleep can also reflect activity patterns of place cells that newly emerge in a familiar environment ^24^. These data all suggest that after prolonged sleep, some of the patterns can reflect novel patterns seen after sleep in novel or familiar environments.

Can the reactivation drift in our data be related to preplay or the representational drift of place representations seen across familiar environments? Our data analysis used a pHMM model to identify distinct cell assemblies, which only took into account the correlated temporal firing structure of the recorded pyramidal cell population. However, the waking activation of pHHM states exhibited spatial selectivity, indicating that these corresponded to distinct assemblies of place cells representing different locations. Accordingly, the activation of recall states in the prior rest session suggests that some of the updated place cell assemblies of the recall session already emerged in prior sleep. We could further confirm this effect by using a Bayesian place-decoding reactivation analysis (S2g-h).

Considering that we observed this effect in a familiar environment (albeit during novel spatial goal learning), our findings may also be related to the representational drift observed across hippocampal place cells over repeated exposure to familiar environments ^21,22,25^. However, our experimental conditions were different from the preplay studies in which an entirely novel environment was introduced. In our case, the animal learned a novel set of reward locations in a familiar environment and the prolonged rest period separated the next day’s recall. This condition was actually similar to a recent work in which extra-stable recording in two-photon imaging was performed daily while animals learned and recalled a reward zone on a running belt ^26^. In this study, 35% of place cells maintained similar spatial representations across days. This proportion is somewhat similar (25%) to our stable cell groups that maintained stable firing rates across learning and recall. The spatial population vectors of the stable cells strongly correlated (r=0.9) across learning and recall sessions, demonstrating that, indeed, this group maintained stable spatial representations for longer than a day. However, the other two groups exhibited lower but still relatively strong (decreasing: 0.5 and increasing 0.7) population vector correlations, indicating that some of these cells also maintained similar place fields while others cells may have remapped. Also, a significant portion of the rate-changing cells remained active during the entire recording (top 50% exhibited >0.3 Hz). However, the relatively higher population vector correlations and sustained activity at a lower firing rate suggest that many of the rate-changing cells may have undergone rate remapping.

### Stable and rate-changing pyramidal cell groups

Several studies demonstrated that hippocampal pyramidal cells are not uniform and can be subdivided into groups based on their anatomical or physiological properties. Initially, CA1 pyramidal cells have been differentiated according to their calbindin expression ^27–32^. Later work showed that CA1 pyramidal cells could be subdivided according to their oscillatory firing properties, firing rate, and burst propensities ^33^. In our classification, we compared the firing rate distribution of cells across learning and recall sessions to separate them into stable-rate persistent cells and rate-changing groups that either increased or decreased their rate from learning to recall. We showed that persistent cells alone did not exhibit reactivation drift, while the rate-changing groups did. Interestingly, about half of the pyramidal cells belonged to the rate-reducing group, which cells also exhibited a rate reduction whenever the newly emerging recall pHMM states were activated in sleep. This is consistent with the homeostatic rate reduction role of sleep and further suggests a more sparse place representation to emerge after sleep ^34^. We also observed a negative correlation between firing rate changes across REM and NREM sleep, which is in line with the opposing role of these states in firing rate regulation ^35^.

In relation to replay and preplay, plastic and rigid cell groups have been differentiated: rigid cells participated in the preplay of firing sequences, while plastic cells refined replay sequences in subsequent sleep without contributing to preplay ^19^. Overall, in that study, the rigid cells exhibited higher firing rates and reduced spatial selectivity compared to the plastic group. As mentioned above, reminiscent of preplay, we observed the emergence of updated recall assembly patterns before, during the rest session. One might expect that the rigid group identified during preplay is equivalent to the persistent group in our classification that maintained unaltered firing during sleep; hence, many of these could belong to ‘pre-wired’ assemblies. However, not only the activity patterns of persistent cells could be seen in prior sleep but those of the increasing cells as well. Moreover, the rate-changing cells exhibited more similar firing characteristics to the rigid cell groups seen in preplay: they had overall higher firing rates than the persistent cells, and when they fired at lower rates, their spatial selectivity was also reduced compared to persistent cells. However, we expect that the rate-changing groups represent plastic cells, considering that they reorganized their firing relationships to other cells during sleep. As we point it out later, a possible explanation for this could be that the rigid cell group seen in preplay were in fact represented not rigid but plastic assemblies that emerged in sleep.

### Network mechanism of reactivation drift

As we saw above, the persistent cells exhibited stable reactivation and spatial coding for the entire duration of the recordings. These cells probably overlap with those place cells that, in calcium imaging experiments, maintained stable place fields across days ^26,36^. These cells can enable continuity and provide a reference frame for the transformation of reactivation patterns. At the same time, we expect that the update of reactivated spatial assemblies would require plasticity that enables new members of the increasing group to associate with existing members of the persistent group and the decreasing group to decouple from them.

SWRs themselves may provide the means for plastic changes to enhance the assembly association of new members ^37^. Previously it was shown that if a CA1 cell is repeatedly excited during a SWR, that cell will increase its SWR-associated firing after the pairing ^38^. We expect that assembly-specific SWR pairing will increase the cell’s firing only when that paired assembly is activated during SWRs considering that SWR-like synchronization enhances plasticity of cells that are co-active ^39,40^. The drift towards the updated assemblies primarily occurred during NREM sleep in the presence of SWRs. Therefore, we speculate that the association of new members to an assembly may be initiated by a random process in which some non-member cells may start to fire with a specific assembly. If that cell has a lower threshold for plasticity, even a few random pairing events may initiate a self-reinforcing process in which pairing will lead to increased activation probability with an assembly, which will lead to further pairing and a further increase in assembly association.

In parallel with the opposing role of REM sleep in reactivation drift, plastic changes during REM sleep may favor assembly stabilization instead of assembly drift. Plastic changes that are thought to enhance the stabilization of newly formed place fields in waking theta oscillations may operate in a similar manner during REM sleep as well. These may also enable the strengthening of existing assemblies. This process may also extend to updated assemblies that emerge in previous NREM sleep and enable them to be maintained in subsequent waking activity as well. Given that theta oscillations can facilitate both LTP and LTD ^41,42^, not only associations of new members could be enhanced during REM sleep, but the decoupling of other cells from the rate-decreasing group could also occur.

A number of additional factors may also impact different reactivation dynamics and associated circuit functions during REM and NREM epochs ^43^. Many of these could be attributed to different physiological states due to the different levels of neuromodulation provided by acetylcholine or other non-specific neurotransmitters, including serotonin ^44,45^. Moreover, sleep stages are also different in terms of large-scale network interactions across the entire brain ^5,45–47^. One could speculate that enhanced out-of-assembly activation during SWR events might have originated from bi-directional interactions of the hippocampus with other cortical centers^48^. In the same way, the strengthening of existing assemblies during REM periods could be allowed by the relative isolation of the hippocampus from external sources of information, which is thought to be a hallmark of REM periods of sleep ^49–51^.

The process we outlined may point to a general sleep mechanism that is not restricted to the case of updated spatial maps in familiar environments or similar spatial learning contexts seen across days. Similar processes may help the rapid initial formation of novel cognitive maps ^52^ as well and may even enable preplay ^18–20^. Ultimately, updated assemblies that emerge during prolonged sleep/rest periods could be used to provide the initial building block for cognitive maps that emerge during the first exposure to a novel environment. Hence, preplay may not exclusively represent prewired assemblies but also assemblies that dynamically emerge during long periods of rest and sleep. Unlike fixed assemblies, this can enable a wide variety of assembly patterns to emerge in sleep, ensuring the emergence of diverse assemblies representing distinct cognitive maps. Similar assembly reorganization involving the gradual morphing of assemblies in sleep could also take place in other brain areas.

## Acknowledgments

We thank Andrea Cumpelik and Freya Ólafsdóttir for comments on an earlier version of the manuscript. This work was supported by the European Research Council (281511) and Austrian Science Fund (FWF I3713).

## Author Contributions

F.S., P.B and J.C. designed the study. P.B conducted the experiments and F.S. and L.B. analyzed the data. F.S., L.B. and J.C. wrote the manuscript.

## Supplementary figures

**Supplementary Figure 1:**
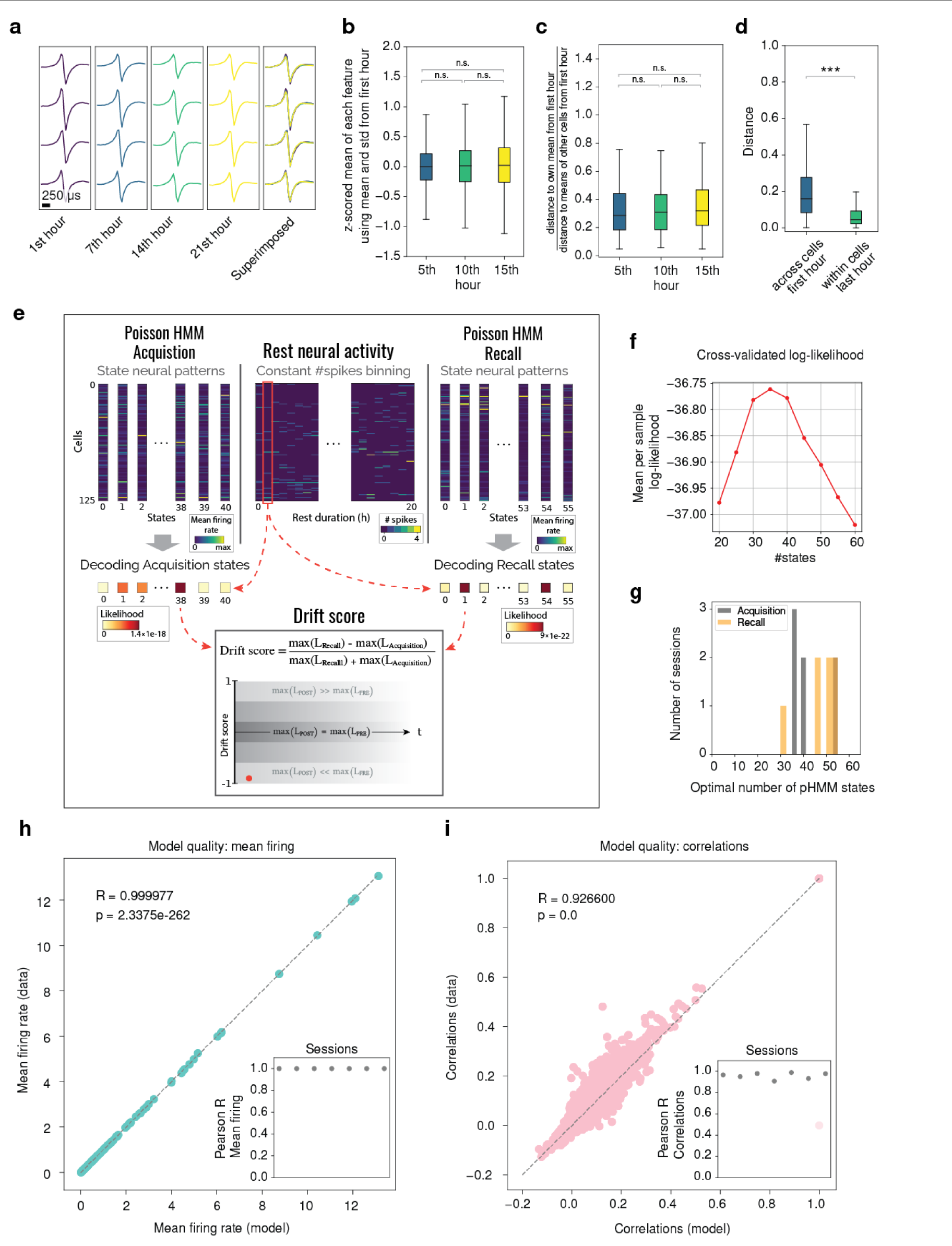
**(a)** Example spike waveform of one hippocampal principal cell unit (over tetrode channels) for the ∼20 hs experiment duration. **(b)** Temporal stability of the first 12 clustering features. For each feature, the mean was computed for the corresponding interval (5th, 10th, 15th hour) and z-scored using the mean and SD of the first hour (for all sessions p > 0.4, two-sided Mann-Whitney U test). **(c)** Distance (1 - Pearson R) between the mean cluster (first 12 clustering features) at 5, 10 and 15 hours relative to the same cluster mean of the first hour and the other cluster means of the first hour. Ratio of within vs. across cluster distance is shown for one example session (for all sessions p>0.15, two-sided Mann-Whitney U test). **(d)** Across cells distance: mean cluster distance (1-Pearson R, 12 clustering features) across cells for the first hour. Within cells distance: distance between the cell’s mean from the first and the last hour of the recording for one example session (for all sessions p < 0.001, two-sided Mann-Whitney U test). **(e)** Decoding neural activity during sleep. Two separate pHMMs were fit to the waking neural data from acquisition and recall, respectively. Sleep neural data is binned using a constant number of 12 spikes per bin. The data likelihood for each bin is computed using acquisition and recall states from the pHMMs. The drift score is calculated using the maximum likelihoods from acquisition and recall. **(f)** Cross-validated log-likelihood to identify optimal number of hidden states (example session). **(g)** Optimal number of states for acquisition and recall for all sessions. **(h)** Mean firing rates of real data versus mean firing rates of data sampled from the pHMM for one example session (R = 1.0, p =2.3375e-262). Inset: Pearson R values for all sessions (all R > 0.99088). **(i)** Correlation of instantaneous firing rate of cell pairs of real data versus sampled data from the pHMM for one example session (R = 0.967, p = 0.0). Inset: Pearson R values for all sessions (all R > 0.85709).

**Supplementary Figure 2:**
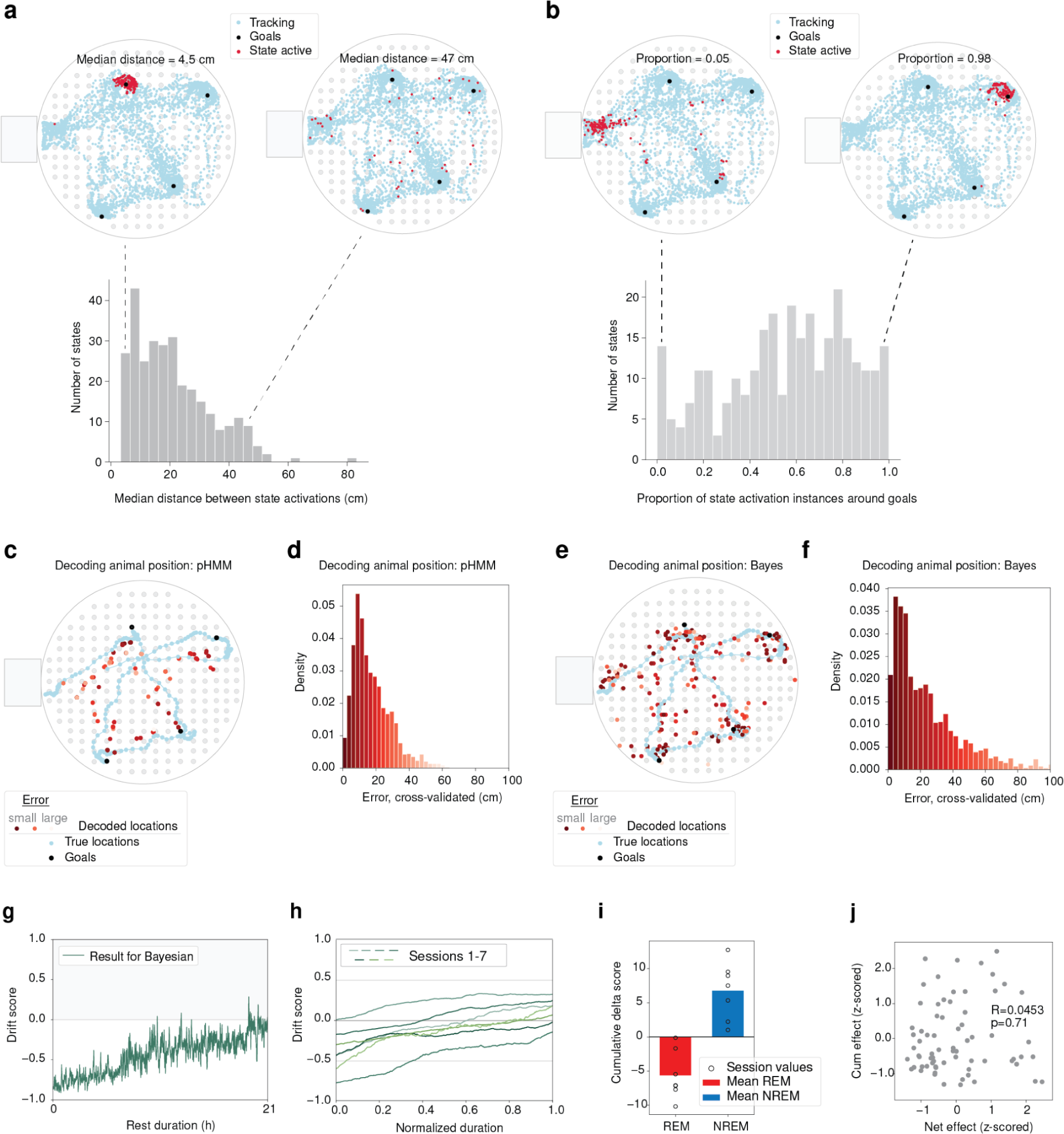
**(a,b)** Top: examples showing the cheese board locations where a given pHMM state was decoded during acquisition. Bottom: (a) distribution of median distances of decoded locations for each pHMM state for all sessions. **(b)** Distribution of goal selectivity of pHMM states measured as the proportion of instances when a pHMM state was active near (<10cm) a goal for all sessions. **(c)** True and decoded locations using the pHMM states for decoding (see Methods) for one example trial. **(d)** The distribution of cross-validated dDecoding errors using the pHMM decoding approach for all sessions. **(e-f)** Same as (c-d) but using Bayesian decoding. **(g)** Drift score using Bayesian decoding (one example session). **(h)** Drift score using Bayesian decoding for all sessions. **(i)** Cumulative ΔDrift score for REM and NREM for all sessions using Bayesian decoding of sleep activity. **(j)** Correlation between cumulative effect (sum of absolute ΔDrift score values per epoch) and net effect (difference in ΔDrift score between the end and the beginning of each epoch) for all sleep epochs (R=0.0453, p=0.71, data from all sessions).

**Supplementary Figure 3:**
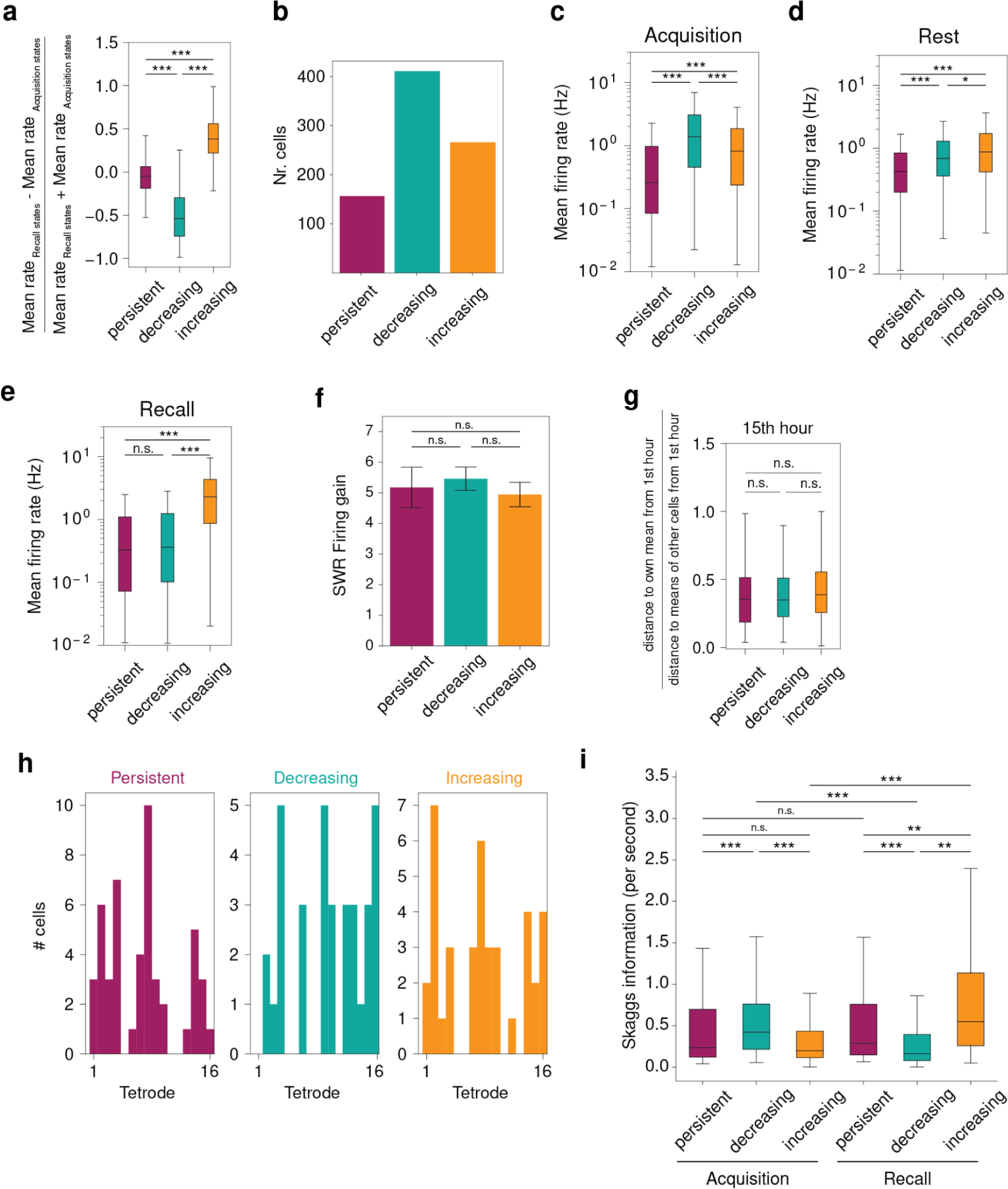
**(a)** Normalized difference in acquisition and recall rates extracted from pHMMs state vectors (Figure S1e) for persistent, decreasing and increasing cells (*** p < 0.001, two-sided Mann-Whitney U test, Bonferroni correction). **(b)** Number of persistent, decreasing and increasing cells for all sessions. **(c-e)** Mean firing rates during acquisition **(c)**, rest **(d)** and recall **(e)** sessions (*** p < 0.001* p < 0.05, n.s. p > 0.05, two-sided Mann-Whitney U test, Bonferroni correction). **(f)** SWR firing rate gain (mean±SEM, ratio of baseline to SWR peak firing rate) for persistent, decreasing and increasing cells (all p>0.29, two-sided Mann-Whitney U test). **(g)** Cluster stability of different cell groups. Distance (1 - Pearson R) between the mean cluster (first 12 clustering features) between first and fifteenth hour. Ratio of within vs. across cluster distance is shown. Data from all sessions (all p > 0.05, two-sided Mann-Whitney U test). **(h)** Number of persistent, decreasing and increasing cells detected across the different tetrodes of an example recording session. **(i)** Skaggs spatial information for persistent, decreasing and increasing cells during Acquisition and Recall (*** p < 0.001, ** p < 0.01, n.s. p > 0.05, two-sided Mann-Whitney U test, Bonferroni correction).

**Supplementary Figure 4:**
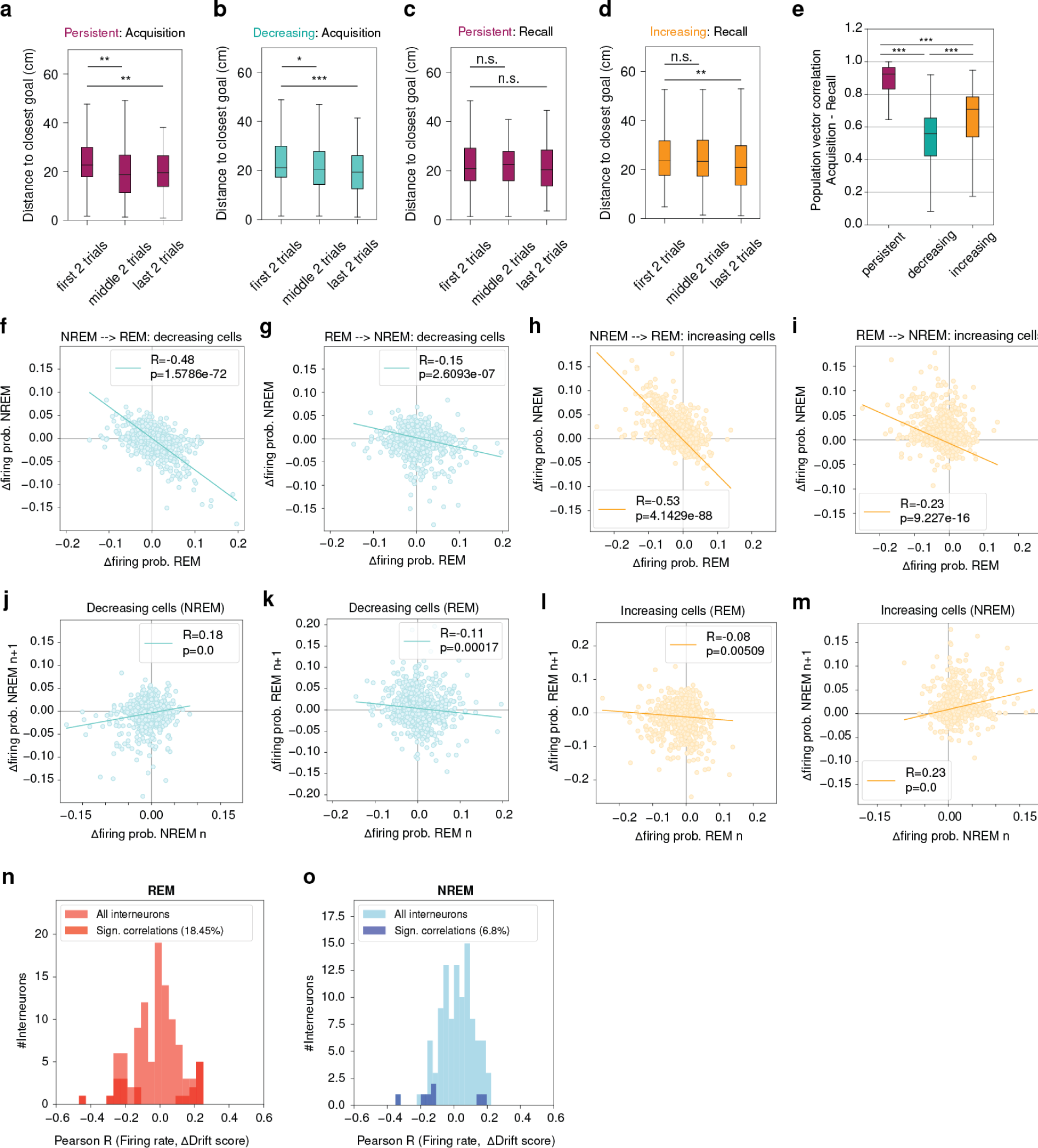
**(a-b)** Distances between place field peak firing locations and goal locations during acquisition for persistent **(a)** and decreasing **(b)** cells (* p < 0.05, ** p < 0.01, *** p < 0.001, two-sided Mann-Whitney U test, Bonferroni correction). Data from all sessions. **(c-d)** Same as (a-b), but persistent and increasing cells are shown during recall (** p < 0.01, n.s. p > 0.05, two-sided Mann-Whitney U test, Bonferroni correction). **(e)** Population vector correlations between acquisition and recall for persistent, decreasing and increasing cells (*** p < 0.001, two-sided Mann-Whitney U test, Bonferroni correction). **(f)-(i)** Correlation between changes in firing probability in neighboring NREM and REM periods for **(f)-(g)** decreasing and **(h)-(i)** increasing cells considering the order of subsequent sleep type periods (R < −0.47, p < 1.579e-71 for NREM followed by REM and R < −0.14, p < 2.609e-6 for REM followed by NREM). **(j)-(m)** Correlation between firing probability changes in subsequent sleep epochs of the same type for **(j)-(k)** decreasing and **(l)-(m)** increasing cells (R < 0.24 for NREM, R < −0.07 for REM). **(n)** Distribution of Pearson R values between interneuron firing rate and ΔDrift score for all interneurons from all sessions for REM epochs. 18.45% of Pearson R values were statistically significant (p<0.05, binomial test p < 0.001). **(o)** Same as (a) but for NREM epochs. 6.8% of Pearson R values were statistically significant (p<0.05, binomial test p > 0.05).

## Methods

### Animals and surgery

Animals were implanted bilaterally with microdrives housing 32 (2×16) independently movable tetrodes targeting the dorsal CA1 region of the hippocampus. Each tetrode was fabricated out of four 10 μm tungsten wires (H-Formvar insulation with Butyral bond coat California Fine Wire Company, Grover Beach, CA) that were twisted and then heated to bind them into a single bundle. The tips of the tetrodes were then gold-plated to reduce the impedance to 200-400 kΩ. During surgery, the animal was under deep anesthesia using isoflurane (0.5%–3% MAC), oxygen (1-2l/min), and an initial injection of buprenorphine (0.1mg/kg). Two rectangular craniotomies were drilled centered above CA1 and positioned relative to bregma (centered at AP = −3.2; ML = ±1.6), the dura mater removed and the electrode bundles implanted into the superficial layers of the neocortex, after which both the exposed cortex and the electrode shanks were sealed with paraffin wax. Five to six anchoring screws were fixed onto the skull and two ground screws (M1.4) were positioned above the cerebellum. After removal of the *dura*, the tetrodes were initially implanted at a depth of 1-1.5 mm relative to the brain surface. Finally, the microdrive was anchored to the skull and screws with dental cement (R*efobacin Bone Cement R*, Biomet, IN, USA). Two hours before the end of the surgery the animal was given the analgesic Metacam (5mg/kg). After a one-week recovery period, tetrodes were gradually moved into the dorsal CA1 cell layer (*stratum pyramidale*). After completion of the experiments, the rats were deeply anesthetized and perfused through the heart with 0.9% saline solution followed by a 4% buffered formalin phosphate solution for the histological verification of the electrode tracks.

### Data Acquisition, Training and Behavior

The animals were housed individually in a separate room under a 12h light/12h dark cycle. Following the postoperative recovery period, rats were reduced to and maintained at 85% of their age-matched preoperative weight. Water was available *ad libitum*. Each animal was handled and familiarized with the recording room and with the general procedures of data acquisition. Behavioral training was performed after electrodes implantation during days when the electrodes were moved towards the hippocampus, but before they reached the hippocampus. Overall recordings were performed in three rats that were trained to perform the seek of the hidden rewards task on the cheeseboard maze ^8,53^ and come back to the start-box. In order to achieve this, random groups of visible food pellets (MLab rodent pellet 20 mg, TestDiet) were spread out on the surface of the cheeseboard maze while the rat was inside the start box. Then we opened the door and left the animal freely foraging the entire maze and once the animal returned to the start box we closed the start-box door. With the help of this training protocol we could shape our animals behavior to automatically explore the entire maze and return to the start-box. Despite the automatic behavior we could ensure that under experiments rats had no or limited experience in performing the cheese-board maze task during the time of the recordings. Each daily experiment consisted of a sequence of nine recording sessions in the following order: a free exploration session on a familiar environment, half hour immobility/sleep rest session in the animal’s own cage, free exploration on the cheese-board, an immobility/sleep rest session (own cage), a learning session (4 randomly selected invisible locations) on the cheese-board, 20 hour continuous monitoring in the cage, recall learning session with the same bait locations, an immobility/sleep rest session (own cage), post probe on the cheese-board (free exploration, unrewarded) and another free exploration session on the same familiar environment as the first one. During the learning session once animals learned the four invisible goal locations, they performed only 5 additional trials to master the task. Following the ∼20 hour long rest session we tested the recall performance of the animal on the cheeseboard maze in 50 further rewarded trials with the same bait locations which were learned 20 hours before.

To be able to record extracellular electric signals continuously over a long period of time we adapted to our commonly built 3D printed microdrive a new, high-fidelity 64-(two animals) and 128-channel (one animal) wireless recording system from TBSI (Triangle BioSystems, Durham, NC, W128) to use in our experiments. In animals in which only the 64-channel telemetry system was available, only half of the recorded channels of the 128 channel drive were used. Using this experimental preparation, we recorded cell population activity continuously over 30 hours during learning, long periods of rest where reactivation takes place and memory recall tests in the end. This telemetry system has been able to amplify and transmit wide band (0.7 Hz to 9 kHz) signals above 20 kHz on the analog channels, which were then digitized at 20 kHz.

Two small light-emitting diodes (LEDs) mounted on the preamplifier headstage were used to track the location of the animal via an overhead video camera. The animal’s location was constantly monitored throughout the experiment. We detected the two LEDs with a custom made tracking software (*positrack*, github.com/kevin-allen/positrack) made by Kevin Allen. The video signal has been triggered and tracked continuously with a TTL pulse sent by the camera’s computer on a common analog channel.

### Spike Detection, Sorting and Stability

The spike detection and sorting procedures, and clustering were performed as previously described ^54,55^. Continuously recorded wide-band signals were digitally high-pass filtered (0,8-5 kHz). Action potentials were extracted by first computing power in the 800-9000 Hz range within a sliding window (12.8 ms). Action potentials with a power of > 5 SD from the baseline mean were selected and spike features were then extracted by using *principal components analyses* (PCA). The detected action potentials were segregated into multiple single units by using automatic clustering software (http://klustakwik.sourceforge.net/). These clusters were manually refined by a graphical cluster cutting program. Only units with clear refractory periods in their autocorrelation (<20μs) and well-defined cluster boundaries were used for further analysis. We further confirmed the quality of cluster separation by calculating the Mahalanobis distance between each pair of clusters ^56^. To be able to analyze changes in the firing patterns of neuronal ensembles over time, we had to guarantee that our set of putative cells was sampled from clusters with stable firing over the whole recording. To ensure this we clustered together periods of waking and rest sessions and then we plotted spike features over time by plotting 2-dimensional unit PCA cluster plots across the whole recording in addition to the stability of spike waveforms. With the help of this method we could exclude those spike clusters which overlapped during the course of recordings. To further verify spike cluster stability we used the *t-student stochastic neighbor embedding* (t-sne) dimensionality reduction method: T-sne embeds the n-dimensional extracellular spikes (n = number of features by which each spike is decomposed) into a low dimensional space ^57^. T-sne focuses on ensuring that the local structure remains intact while it ignores the global structure, therefore when we expressed T-sne features over time we could visually exclude those clusters which were unstable during the whole recording due to electrode drifting.

Putative pyramidal cells and putative interneurons in the CA1 region were discriminated by their autocorrelations, firing rate, and waveforms, as previously described ^54^.

### Sleep classification

In recordings, exploratory and immobility or sleep sessions were manually separated off-line as previously described ^54,55^. For each session, the theta/delta ratio was plotted against speed so that the behavioral state could be manually identified. The theta/delta field power ratio was measured in 1,600 ms segments (800 ms steps between measurement windows) with the Thomson’s multitaper method ^58,59^. Waking behavior included periods of locomotion and/or the presence of theta oscillations (visible in the theta/delta ratio), with no more than ∼2.5 s of transient immobility. Rest epochs were selected when both the speed and theta-delta ratio dropped below a pre-set threshold (speed: <4cm/s, theta/delta ratio: <2) for at least 2.4 s. During periods of active waking behavior, theta-oscillatory waves detection was performed as previously described ^54,55^ using the negative peaks of individual theta waves from the filtered trace of the local field potential (5–28 Hz). The band used for the detection was wider than the theta band in order to precisely detect the negative peaks of the theta waves, which otherwise would have smoothed out in using a narrow theta band. Sleep segments have been identified by longer periods of immobility (because of the lengths of our recording at least 10 min) and the clear presence of REM-theta and slow-wave field oscillations.

### Sharp wave ripple detection

For the detection of SWRs, local field potentials were band-pass filtered (150–250 Hz), and a reference signal (to ensure the lack of ripple activity we left a tetrode above the hippocampus as a reference) was subtracted to eliminate the so called common noise (muscle artifacts due to scratching, twitching etc.). The power (root mean square) of the filtered signal was calculated for each electrode and summed across electrodes considered to be in the CA1 *stratum pyramidale*. The threshold for SWR detection was set to 7 SD above the background mean. The SWRs detection threshold was always set in the first sleep session, but longer (at least 35 min) as it was described earlier and the same threshold was used for all other sessions ^54,55^.

### Quantification of stability of clustering features over time

To assess the temporal stability of waveforms, we used the first 12 clustering features. For each feature, the mean was computed for the corresponding interval (5th, 10th, 15th hour) and z-scored using the mean and std of the first hour.

### Cell separation over time

We estimated the separation of cells by using the first 12 clustering features and three different approaches. First, for each cell, we divided the distance (1 - Pearson R) between the mean value of each feature during the nth hour and the cell’s own mean feature values from the first hour by the distance between the mean feature values during the nth hour and the other cells’ feature means from the first hour. Using this measure we investigated whether a cell’s waveform at a later stage during the experiment is more similar to its own waveform from the first hour than to first hour waveforms from other cells. Second, for each cell, we divided the distance (1 - Pearson R) between the mean feature values during the nth hour and the cell’s own mean values from the first hour by the distance between the other cells’ feature means during the nth hour and the cell’s own feature means from the first hour. This measure estimates whether other cells’ waveforms at later stages during the experiment are more similar to the cell’s waveform from the first hour than its own waveforms at later stages. Third, we compared the across cell distance with the within cell distance. For the across cell distance, the distance between the mean feature values per cell of the first hour and the other cells’ feature means from the first hour was calculated. The within distance corresponds to the distance between the cell’s feature averages from the first hour and the same cell’s feature averages of the last hour of the experiment.

### Clustering feature stability

For each cell we calculated the mean and std for each of the first 10 PCA clustering features using data from the first hour. We then computed the mean per clustering feature of each cell at different time intervals (7^th^, 15^th^ and 21nd hour) of the experiment. In order to test whether clustering features drift away from the initial values, we z-scored the means during different time intervals using the mean and std from the first hour.

### Measuring excess path during acquisition and recall

We assessed the animal’s ability to learn and recall goal locations on the cheeseboard by computing the excess path: once the rat had left the start box, we measured the length of the path the animal took to reach any of the four goals (animal position within 10 cm radius around goal location). Next, we detected when the animal left the goal again and measured the path length to the next goal. This procedure was repeated for the remaining two goals. We then calculated the optimal paths as straight lines between either start location and the first goal or between subsequently visited goals. Each taken path length was then divided by the optimal path length to yield the excess path as a multiple of the optimal path.

### Poisson hidden markov model (pHMM) and model fitting

We trained two separate hidden Markov models with Poisson emissions (pHMM) on the neural data obtained during the cheeseboard task before (acquisition) and after (recall) the long sleep. Only data from running periods (speed > 5cm/s) was used. The acquisition data length was matched in terms of duration to the recall data to have the same training data length for acquisition and recall. Then, the neural data was binned using temporal bins of 100 ms length.

The pHMM model assumes the temporal evolution of an unobserved discrete state as described in ^60^. In short, the probability of observing an ensemble *E_N_* of N independently firing neurons at time t for mode *I* can be modeled as:

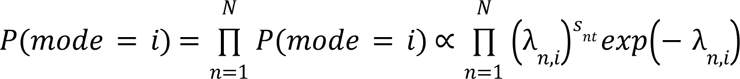

where s_nt_ is the firing rate of neuron n at time t. The firing rate is modeled according to a Poisson process with a mean firing rate λ_n,i_ defined by the unobserved discrete state I.

The transition probabilities between M unobserved modes is captured by the M x M transition matrix **A.** The hyperparameter M defining the number of modes of the model was determined using the cross-validated maximum likelihood (Supplementary Figure 2). All model parameters were computed using the EM-algorithm.

### Rate map generation for Bayesian decoding

For each session and cell one rate map for acquisition and one rate map for recall was computed.

In order to reconstruct the two-dimensional spatial distribution of each cell’s firing probability, we used a maximum entropy model inference paradigm ^61^. Only running periods were considered by applying a 5 cm/s speed filter. The cell activity was binned using time windows of 10 ms. We assigned a binary variable S_i_(t) to each neuron to denote the presence or absence (+1/−1) of spikes emitted by neuron I within time bin t. Including the dependence of each neuron’s state on the previous state (t-1) of the population, the maximum entropy distribution over the state S_i_(t) of neuron I at time t is

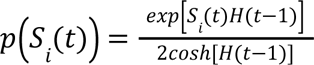

with H(t) being the time-varying covariate representing the external field in statistical physics. Equation (1) describes a Generalized Linear Model (GLM) with the interaction kernel extending to one time step in the past.

By maximizing the log-likelihood with respect to H(t) we found the most likely values of H(t) to generate the observed data:

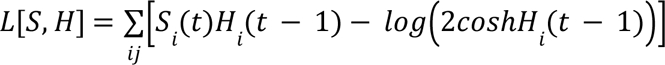

For the spatial input we assumed the sum of two-dimensional Gaussian basis functions centered on an evenly spaced M x M square lattice that spanned the cheeseboard. The spatial field of cell I at time t can then be computed in the following way:

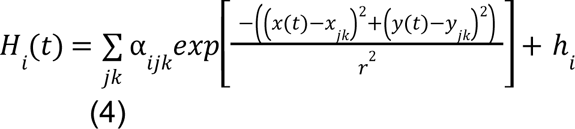

where *h_i_* is the baseline activity of cell I, (x_jk_, y_jk_) are the centers of the Gaussians and *r* is the standard deviation of basis functions. An accurate representation of the cell activity in space was computed by inferring parameters *α_ijk_* of the linear combinations of Gaussian basis functions. The resulting map was then partitioned into bins of 4 cm.

### Decoding sleep activity using pHMM

In order to compensate for differences in temporal dynamics between REM and NREM sleep, we binned the sleep data using bins with a constant number of 12 spikes. Since the awake pHMM models were trained on temporal bins of 100 ms we computed a scaling factor between awake and sleep neural activity to match the two. First, we calculated the mean number of spikes occurring within 100 ms time bins during awake behavior n_awake,100ms_. The scaling factor γ_phmm_ is defined as:

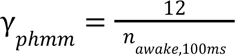

The likelihood of the sleep activity at time *t* given the discrete pHMM state I is computed as follows:

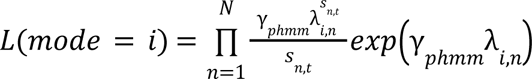

where *S_t_* is the sleep activity of *N* neurons at time *t, s_n,t_* is the number of spikes of neuron *n* at time *t* and *λ_i,n_* is the mean firing rate of that neuron in state *I*. Notice that for our decoding procedure the transition probabilities across states were not considered. To assess which state was most likely reactivated at time *t* during sleep the state with the maximum log-likelihood was selected.

### Decoding sleep activity using Bayesian decoding

Equivalent to the pHMM decoding approach the sleep data was binned using bins with a constant number of 12 spikes. In the case of Bayesian decoding the acquisition and recall rate maps were computed using 10 ms time bins. Therefore, the scaling factor γ_bayesian_ is computed using the mean number of spikes occurring within 10 ms time bins n_awake,10ms_ during awake behavior:

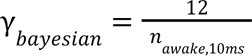

The likelihood of the sleep activity at time *t* given the spatial bin x is given by:

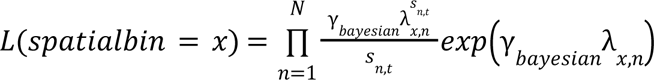

where *S_t_* is the sleep activity of *N* neurons at time *t, s_n,t_* is the number of spikes of neuron *n* at time *t* and *λ_x,n_* is the mean firing rate of that neuron in spatial bin *x*. To reconstruct the spatial bin that was most likely reactivated at time *t*, the spatial bin with the maximum log-likelihood was selected.

### Quantifying drift during sleep

For each time point *t* in sleep, we calculated the Drift score in the following way:

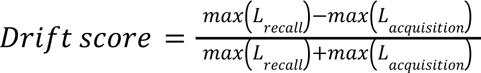

with *L_recall_* and *L_acquisition_* being the maximum likelihoods across all states or spatial bins for the acquisition or recall models (pHMM or rate maps) at time *t*, respectively. The resulting Drift score smoothed across time.

#### First vs. second half of sleep

In order to validate whether the drift is more prominent during the first half of sleep, we computed the delta of the similarity ratio for the first and second half of sleep separately. Then, we tested whether the ratio between the delta of the first half and the delta of the second half was greater than 1 using the student T-test.

### Net effect vs. cumulative effect of drift

In order to assess the amount of memory drift with respect to different timescales, we compared the net effect and cumulative effect. For the net effect, we computed the difference in drift score between the beginning and the end of the rest period. On the other side, the cumulative effect was computed by summing the absolute values of the drift score throughout the rest period.

### Opposing effect on drift in NREM and REM

NREM and REM periods were identified as described above. Neighboring sleep epochs of the same type were merged to obtain a set of alternating NREM/REM epochs. For each epoch, the change in similarity ratio was computed by subtracting the first value of the epoch from the last value of the epoch.

### Persistent and unstable subsets

Persistent and unstable cell subsets were identified based on changes in their firing rate distributions from acquisition to recall. For each cell the distributions of firing rates for acquisition and recall were computed separately. If the acquisition distribution was significantly greater than the recall distribution (one-sided MWU, p<0.01), the cell was assigned to the decreasing subset. If, on the other hand, the recall distribution was significantly greater than the acquisition distribution (one-sided MWU, p<0.01), the cell was labeled as increasing. All other cells which did not show a significant difference in their firing rate distributions from acquisition to recall made up the persistent subset.

### Drift using different subsets of cells

To evaluate the effect of using only a subset of cells for our sleep decoding procedure on the observed drift we proceeded as follows. We removed all cells not contained in the subset from our constant number spike bins and computed the Drift score using the maximum likelihoods from the acquisition and recall states. Then, the Drift score for the entire sleep duration was split into four parts of equal length. For each part, a line was fit and the resulting slope calculated. The mean slope of the four parts was then compared to the equivalent value of the Drift score using the entire cell population.

### Firing probability changes in NREM and REM

REM and NREM sleep epochs were identified as described above. Thereafter, for each subset of persistent, increasing and decreasing cells we computed the change in firing probability from the beginning to the end of each epoch. The firing probability is defined as the number of spikes contributed by the subset to the total number of spikes per constant spike bin. We computed the change by subtracting the firing probability of the first bin of the epoch from the value of the last bin of the epoch. Only epochs with significant changes in firing probability were considered.

### Measuring spatial information

To assess the spatial information of single cells we computed the sparsity and spatial information per second as previously described ^62^. The spatial information per second was computed using the following equation:

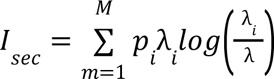

where pi and *λ_i_* are the probability of occupying and the firing rate of bin I, respectively. Parameter *λ* describes the mean firing rate of the cell in the environment.

### Decoding positions using neural activity during behavior

#### Bayesian decoding

We applied standard Bayesian decoding using 1 cm spatial bins. First, we computed the mean number of spikes λ_n,i_ for each cell n and spatial bin i. Given the spikes S_N_ of N neurons at time t, we computed the likelihood of being in bin I using the following equation:

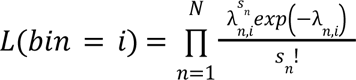

with s_n_ being the number of spikes of neuron *n* at time t. The spatial bin with the highest likelihood represented the decoded location.

#### Decoding position using pHMM

Using the entire neural data from acquisition and the trained pHMM, we inferred the most likely state sequence using the Viterbi algorithm. By matching the sequence of states with the tracking data of the animal we identified a mean spatial location for each state.

Given the activity of N neurons at time t, we computed the normalized likelihood for each state of our pHMM. The decoded location was then calculated by weighing the mean location of each state with its normalized likelihood and computing the average position across all states.

### Computing mean firing rates

Acquisition, sleep and recall were split into 5 min. chunks to computed mean and maximum firing rates of the different cell subsets.

### Distance between peak firing and closest goal

For each cell, we computed its rate map and determined the location on the maze with the highest firing rate. Next, we calculated the distance between the location with the peak firing rate and the closest goal location.

### Population vector correlations

Neurons were separated into persistent, increasing and decreasing cells according to the procedure defined above. Neural activity per subset was binned using spatial bins of 10 cm2 size yielding one population vector per spatial bin. Spatial bins which were not visited in all relevant behavioral episodes were excluded. Then, Pearson correlations between population vectors of the same spatial bin during the different behavioral episodes were computed.

